# Controlling the Mitochondrial Protonmotive Force with Light to Impact Cellular Stress Resistance

**DOI:** 10.1101/742536

**Authors:** Brandon J. Berry, Adam J. Trewin, Alexander S. Milliken, Aksana Baldzizhar, Andrea M. Amitrano, Minsoo Kim, Andrew P. Wojtovich

## Abstract

Mitochondrial respiration generates an electrochemical proton gradient across the mitochondrial inner membrane called the protonmotive force (PMF) to drive diverse functions and make ATP. Current techniques to manipulate the PMF are limited to its dissipation; there is no precise, reversible method to increase the PMF. To address this issue, we used an optogenetic approach and engineered a mitochondria-targeted light-activated proton pumping protein we called mitochondria-ON (mtON) to selectively increase the PMF. Here, mtON increased the PMF light dose-dependently, supported ATP synthesis, increased resistance to mitochondrial toxins, and modulated energy-sensing behavior in *Caenorhabditis elegans*. Moreover, transient mtON activation during hypoxia prevented the well-characterized adaptive response of hypoxic preconditioning. Our novel optogenetic approach demonstrated that a decreased PMF is both necessary and sufficient for hypoxia-stimulated stress resistance. Our results show that optogenetic manipulation of the PMF is a powerful tool to modulate metabolic and cell signaling outcomes.

## INTRODUCTION

Mitochondria generate an electrochemical proton gradient known as the protonmotive force (PMF). This consists of an electrical charge gradient, or membrane potential (Δψ_m_), and a pH gradient (ΔpH) that drives energy availability and controls diverse physiologic outputs (Garrido, Galluzzi et al., 2006, Rizzuto, De Stefani et al., 2012, Shadel & Horvath, 2015). The PMF is generated by proton pumping respiratory complexes of the electron transport chain (ETC) located in the mitochondrial inner membrane (IM). ETC dysfunction can lead to loss of PMF and a diverse range of pathologies (Berry, Trewin et al., 2018, Dingley, Polyak et al., 2010). For example, because the ETC consumes O_2_ to establish the PMF, the PMF is decreased under pathologic hypoxic conditions. The mechanistic link between acute changes in the PMF and downstream physiologic changes are poorly understood (Chalmers, Saunter et al., 2015, Santo-Domingo, Giacomello et al., 2013), and research is focused on developing new techniques to elucidate these pathways (Chalmers, Caldwell et al., 2012, Glancy, Hartnell et al., 2018).

Stroke is a common pathology in which cells undergo hypoxia and rapid reoxygenation that causes changes in the PMF and compromises mitochondrial functions. Changes in the PMF during hypoxia and reoxygenation influence cell-survival outcomes via mechanisms that are not fully understood (Murphy & Hartley, 2018, Solaini, Baracca et al., 2010, Yang, Mukda et al., 2018, Zhang, Trushin et al., 2016). Selectively increasing the PMF to distinguish cause and effect in hypoxic models is necessary to open new avenues of investigation that may reveal important metabolic changes that occur in stroke.

There are several techniques that allow experimental modulation of the PMF, but most are pharmacologic and therefore irreversible and not cell- or tissue-specific. Herein, we take a novel approach to overcome these barriers to precisely control the PMF by using optogenetics, adapting an approach used recently to dissipate the PMF (Ernst, Xu et al., 2019, Tkatch, Greotti et al., 2017). One family of widely-used optogenetic proteins are bacteriorhodopsin-related light-activated proton pumps. These proteins pump protons across membranes in response to specific wavelengths of light, and are often used to study physiology by modulating electrochemical gradients at the plasma membrane (Chow, Han et al., 2010, Husson, Liewald et al., 2012, Kandori, 2015). Only recently have precise optogenetic techniques been applied to compartmentalized cellular events using light-activated proteins targeted to organelles (Ernst, Xu et al., 2018, Rost, Schneider et al., 2015, Tkatch et al., 2017, Trewin, Bahr et al., 2019, Trewin, Berry et al., 2018a). Rather than using a non-specific cation channel to permeabilize the IM and dissipate the PMF, here we target the light-activated proton pump from the fungal organism *Leptosphaeria maculans* (Chow et al., 2010, Waschuk, Bezerra et al., 2005) to mitochondria and selectively increase the PMF. We call this optogenetic tool mitochondria-ON (mtON) due to its ability to mimic the proton pumping activity of the ETC in response to light, independent of oxygen or substrate availability.

We validated mtON using the well-characterized genetic model organism, *C. elegans* (Butler, Ventura et al., 2010, Dingley et al., 2010, Tsang & Lemire, 2003). Using hypoxia and reoxygenation, we tested the hypothesis that hypoxia adaptation through preconditioning requires a decreased PMF. Our data demonstrate that transient loss of PMF during preconditioning is necessary and sufficient for resistance to hypoxia. By probing the evolutionarily conserved hypoxia adaptation response (Pena, Sherman et al., 2016, Wang, Lim et al., 2019, Wojtovich, Nadtochiy et al., 2012b, Wojtovich, Nadtochiy et al., 2013), we show that tools like mtON allow precise determination of cause and effect in physiologic models.

## RESULTS & DISCUSSION

### Light-activated proton pump mitochondria-ON (mtON) is expressed in mitochondria

Using a ubiquitously-expressed gene promoter (P*eft-3*), we directed expression of a light-activated proton pump to the mitochondrial IM in *C. elegans.* Mitochondrial localization was achieved by fusion of the proton pump to an N-terminal mitochondrial targeting sequence of the IMMT1 protein (Fischer, Igoudjil et al., 2011, John, Shang et al., 2005) in an orientation that allows proton pumping from the mitochondrial matrix towards the intermembrane space to increase the PMF in response to light (Fig. 1A). Using a C-terminal GFP fusion for subcellular visualization, and MitoTracker^TM^ CMXRos, we observed overlap of green and red fluorescence in *C. elegans* tissues, indicating the intended mitochondrial targeting (Fig. 1B & Fig. EV 1). We confirmed the expression of mtON in isolated mitochondrial preparations by immunoblot against GFP and observed a band at the predicted molecular weight of 82 kDa (Fig. 1C).

**Figure 1.**
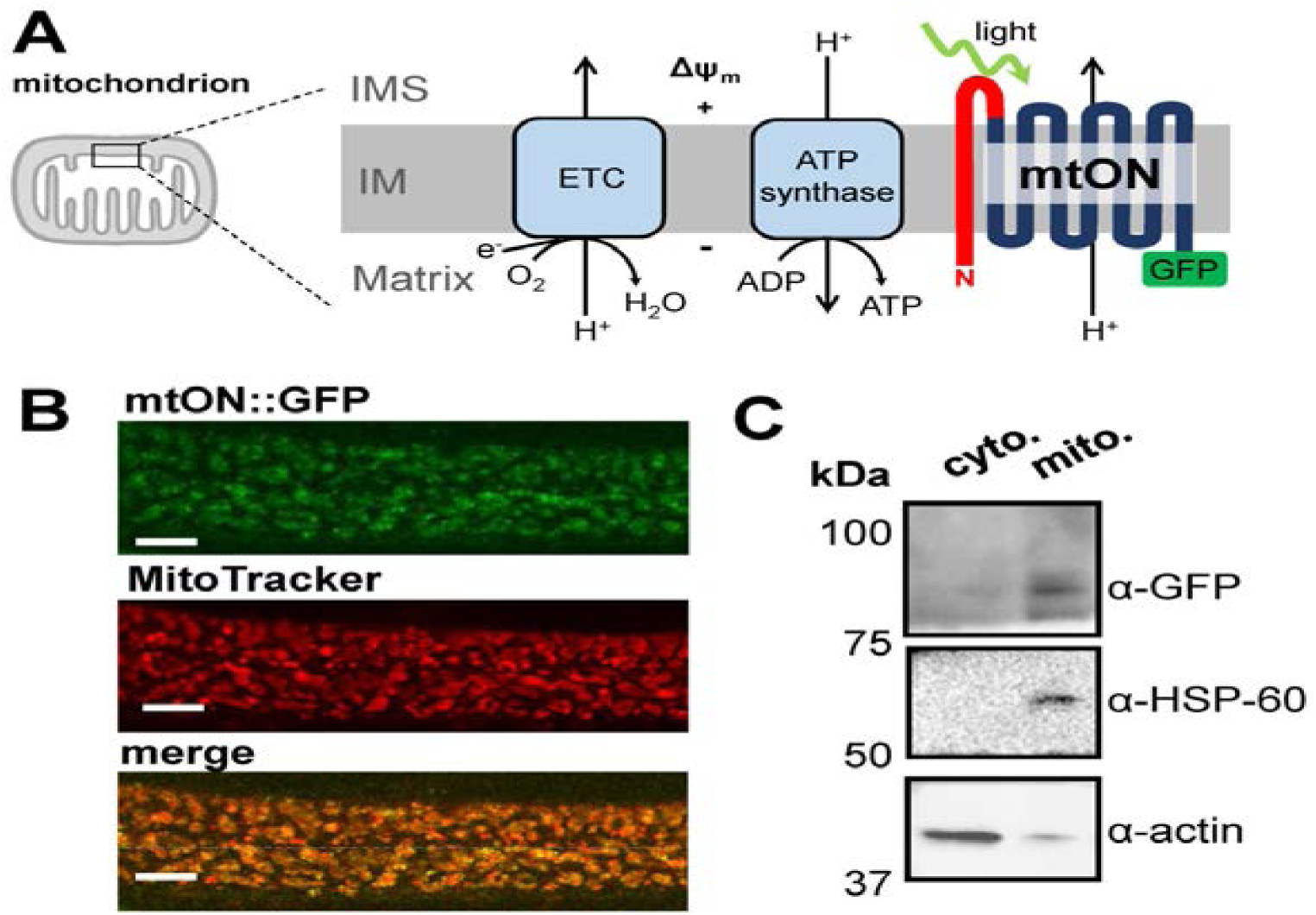
Light-activated proton pump mitochondria-ON (mtON) is expressed in mitochondria. **A** Schematic depicting the targeting strategy to localize mtON to the mitochondrial inner membrane (IM). The electron transport chain (ETC) complexes generate the endogenous mitochondrial PMF by proton pumping, represented by the + and - across the IM. Mitochondrial ATP synthase utilizes the PMF to convert ADP to ATP. The N-terminal mitochondria target sequence from the IMMT1 protein shown in red; GFP shown in green. In response to light, mtON pumps protons from the mitochondrial matrix to the intermembrane space (IMS). **B** Confocal images demonstrate overlap of GFP tagged mtON with MitoTracker^TM^ Red CMXRos stained *C. elegans* hypodermal mitochondria. Scale bar 10 µm. **C** Immunoblot comparing the cytosolic supernatant and the mitochondria-enriched pellet of isolation fractions. GFP-tagged mtON migrates at the predicted molecular weight of 82 kDa accounting for the mitochondria-target sequence, the proton pump, and GFP. mtON is observed only in the mitochondrial fraction compared to marker proteins HSP60 (mitochondria) and actin (cytosol). All blots are from the same lanes on one membrane.

### mtON activation increases the PMF

Proton pumping activity of mtON requires the cofactor all trans-retinal (ATR) (Okazaki & Takagi, 2013, Sumii, Furutani et al., 2005, Waschuk et al., 2005). Because *C. elegans* do not produce ATR endogenously, exogenous supplementation is required for the light-activated proton pump to function (Kandori, 2015, Takahashi & Takagi, 2017). Thus, we were able to control for the expression of mtON (and the included GFP) in functional and non-functional forms, depending on whether ATR was supplemented, in addition to the use of light controls (Fig. EV 2A-D).

To test if mtON was capable of generating a PMF we measured the Δψ_m_ component using the indicator tetramethylrhodamine ethyl ester (TMRE) in isolated *C. elegans* mitochondria. TMRE is a fluorescent lipophilic cation that will accumulate in mitochondria proportionally with the Δψ_m_. At the quenching concentration of TMRE used, fluorescence is low at the Δψ_m_ of intact, energized isolated mitochondria. Upon loss of the Δψ_m_, TMRE redistributes to the extramitochondrial space resulting in increased fluorescence due to dequenching of TMRE (Fig. 2A). Under non-phosphorylating conditions in the absence of added substrate to fuel ETC activity, mtON activation was able to polarize the Δψ_m_ as much as ETC-driven respiration (Fig. 2B) light dose dependently (Fig. EV 3A). These data show that more photons result in increased polarization of the PMF, which is in line with the biophysical studies of the proton pump (Chow et al., 2010). While using the non-functional pump (no ATR), light had no effect on Δψ_m_ (Fig. 2B).

**Figure 2.**
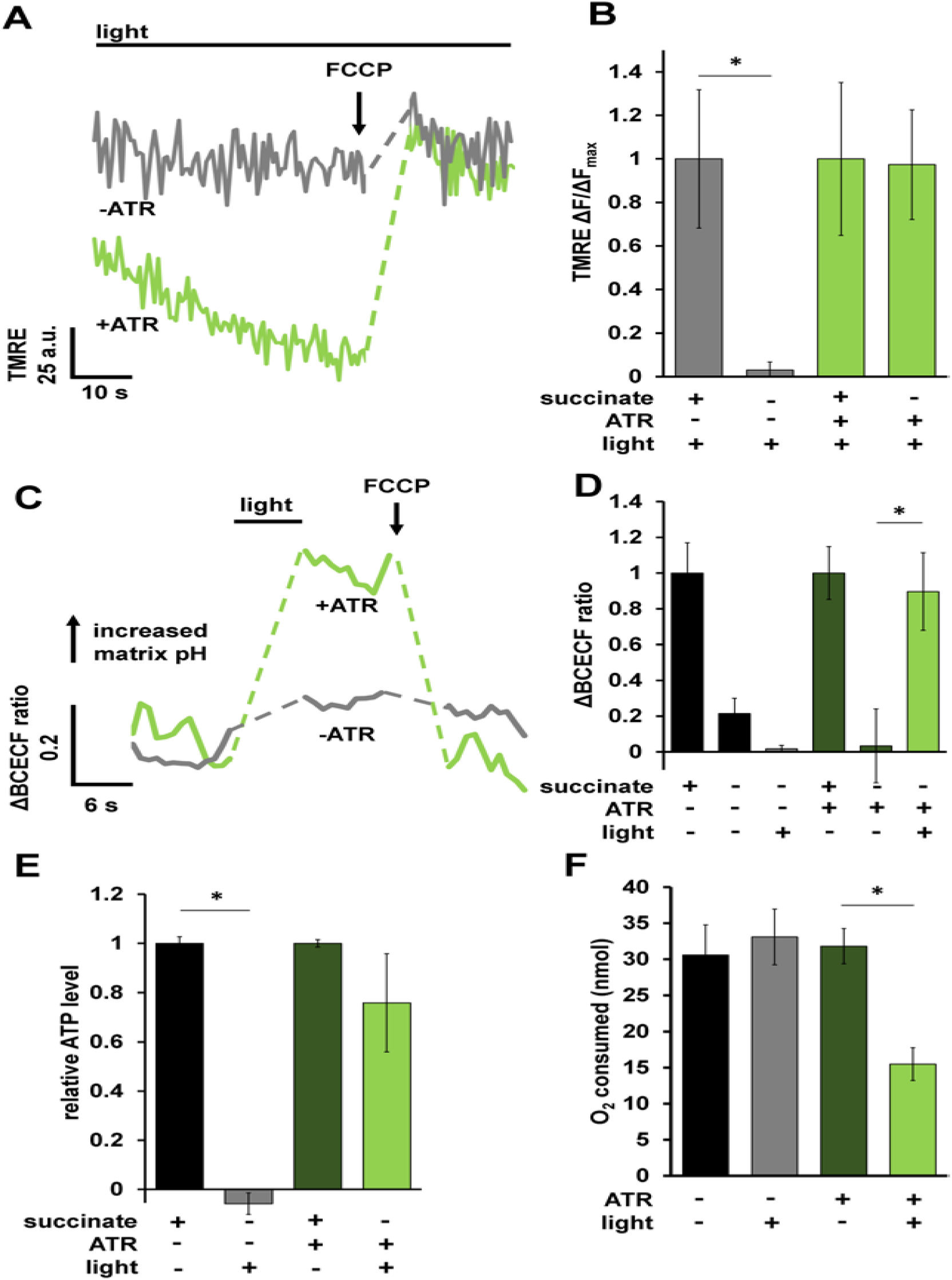
mtON activation increases the PMF. **A** Representative TMRE fluorescence traces in arbitrary units (a.u.) before and after Δψ_m_ dissipation with FCCP. Dashed lines indicate where FCCP was added. +/- ATR traces were performed in the absence of succinate. mtON activation was continuous throughout the traces. Light green trace is from mitochondria with ATR, gray traces are without. **B** Quantification of change in fluorescence (ΔF) normalized to the maximum change in fluorescence given by succinate respiration (ΔF_max_) for each mitochondrial preparation. Data are from the maximum light dose in Fig. EV 3A. Two-way ANOVA with Sidak’s test for multiple comparisons, *p = 0.0469, +ATR succinate vs. +ATR +light p = 0.9978, n = 6 mitochondrial isolations. Bars are means ± SEM. **C** Representative BCECF-AM 490/440 nm ratio trace. Ratio of 545 nm fluorescence intensity at either 440 or 490 nm excitation. Dashed lines are where light or FCCP treatment occurred. Light green trace is from mitochondria with ATR, gray traces are without. **D** Quantification of change in BCECF-AM fluorescence ratio normalized to maximum change given by succinate matrix alkalization. Two-way ANOVA with Sidak’s test for multiple comparisons, −ATR succinate vs. –ATR, +light *p = 0.0212, +ATR succinate vs. +ATR light p = 0.999, +ATR succinate vs. +ATR,-light p = 0.0237, +ATR, +light vs, +ATR, −light p = 0.0474, n = 3 mitochondrial isolations. **E** ATP levels normalized to total ATP synthesis given by succinate respiration. Data from Fig. EV 3B. Succinate data shown for comparison after normalization. Two-way ANOVA with Sidak’s test for multiple comparisons, *p = 0.0011, +ATR succinate vs +ATR light p = 0.5680, n = 3 - 7 assays across two mitochondrial isolations. **F** O_2_ required to consume 50 nmoles of ADP after mtON activation. Data are the maximum illumination from Fig. EV 4D using time-matched dark conditions, One-way ANOVA *p = 0.013 (−ATR, −light vs. +ATR, +light p = 0.010. −ATR, +light vs. +ATR, +light p = 0.013.) −ATR, −light n = 4, rest n = 5 mitochondrial preparations.

When protons are pumped out of mitochondria during respiration the matrix pH increases (Porcelli, Ghelli et al., 2005). Thus, we measured pH changes in the mitochondrial matrix in response to light to test this parameter as another readout of mtON activity. Using a ratiometric pH indicator, BCECF-AM, we found that matrix pH increased in response to mtON activation, mimicking the effect of succinate driven respiration (Fig. 2C&D). mtON-stimulated changes in matrix pH and Δψ_m_ were sensitive to the protonophore FCCP, demonstrating that mtON polarizes the PMF, as measured by both the Δψ_m_ and the ΔpH.

### mtON increases ATP synthesis without respiration

To further test if mtON could increase energetics, we measured ATP levels from isolated mitochondria that were supplied with ADP to phosphorylate, since ATP levels are highly regulated *in vivo* (Balaban, Kantor et al., 1986, Glancy, Hartnell et al., 2015, Viola & Hool, 2017, Wang, Zhang et al., 2017). As expected, mtON activation increased ATP levels (Fig. 2E) light-dose dependently (Fig. EV 3B), similar to the Δψ_m_ results. In mitochondria, the amount of ADP converted to ATP is reliant on O_2_ consumption by the ETC (Brand & Nicholls, 2011). We tested if we could drive ADP conversion to ATP by generating a PMF with mtON, bypassing the requirement for O_2_ consumption. Respiration was consistent across all control conditions (Fig. EV 4A&B), indicating no baseline differences in mitochondrial quality. Activation of mtON decreased the amount of O_2_ required to phosphorylate a given amount of ADP (Fig. 2F & Fig. EV 4C) light dose dependently (Fig. EV 4D), indicating mtON-driven ATP production does not rely on O_2_ consumption. Until now, interventions to increase the PMF have involved fueling the ETC, where here we show mtON generates a PMF independent of O_2_ consumption or metabolic substrates to provide electrons. mtON’s ability to augment the PMF via optic control opens a new avenue for discovery in metabolic research where complex interactions between physiology and bioenergetics often obscure molecular mechanisms.

### mtON increases survival during acute ETC dysfunction

Given that mtON activity decreased reliance on the ETC *in vitro*, we tested if mtON could compensate for acute ETC dysfunction *in vivo*. We exposed *C. elegans* to toxic inhibitors of specific sites within ETC complexes and scored their survival in response to mtON activation. Light alone had no effect on survival after inhibitor exposure in all cases (Fig. 3A-C). For transgenic animals, when exposed to the complex I inhibitor rotenone, mtON was able to improve survival (Fig. 3A). Rotenone toxicity is mediated chiefly through oxidative damage (Ishiguro, Yasuda et al., 2001, Schmeisser, Priebe et al., 2013), and ATR alone exhibited a protective effect under these conditions, possibly due to its antioxidant properties (Lee, Casadesus et al., 2009, Palace, Khaper et al., 1999, Siddikuzzaman & Grace, 2013). This effect was also present in wild type control experiments (Fig. EV 5A). However, the effect of mtON was significantly greater than the ATR effect (Fig. EV 5A, one-way ANOVA, p = 0.0002), with no effect in wild type animals (Fig. EV 5A&B). Activation of mtON also protected animals exposed to antimycin A, a complex III inhibitor (Ishiguro et al., 2001) (Fig. 3B), and azide, a complex IV inhibitor (Fig. 3C). Unlike with rotenone treatment, ATR alone did not mitigate antimycin A or azide toxicity. Overall, these data indicate that mtON can partially overcome inhibited ETC activity in whole organisms.

**Figure 3.**
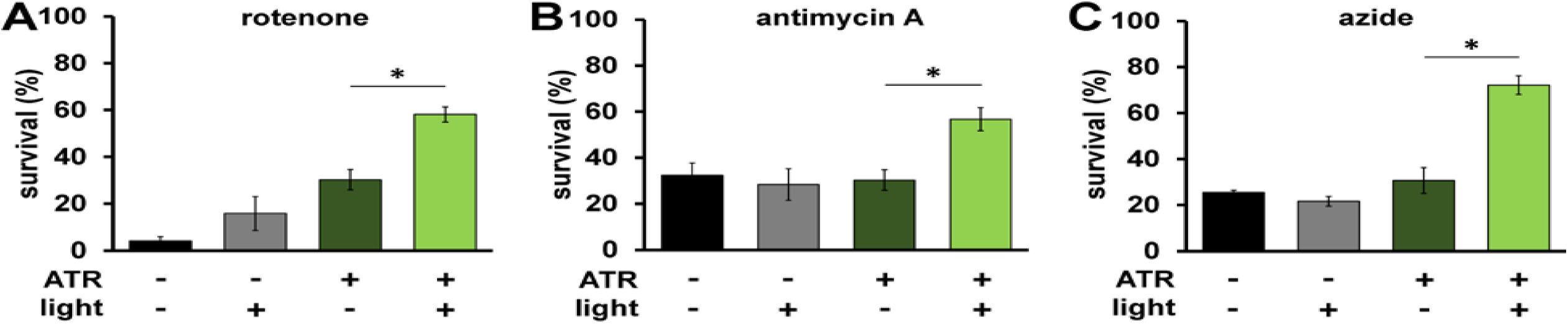
mtON improves survival during acute ETC dysfunction. **A** Day 1 adult animals were exposed to 50 µM rotenone (ETC complex I inhibitor) for 5 hours and survival was scored. Illumination was continuous throughout toxin exposure (see methods). ATR alone was protective (−ATR, −light vs. +ATR, −light p = 0.016.) The effect of mtON activation was greater than the ATR alone effect *p = 0.01. (−ATR, −light vs +ATR, +light p = 0.0002. −ATR, +light vs. +ATR, +light p = 0.0009. n = 3 plates each condition). **B** Animals were exposed to 50 µM antimycin A (complex III inhibitor) and survival was scored 18 hours later. *p = 0.02 (−ATR, −light vs +ATR, +light p = 0.03, –ATR, +light vs. +ATR, +light p = 0.01. n = 5 plates each condition). **C** Animals were exposed to 0.25 M azide (complex IV inhibitor) for 1 hour and scored for survival 1 hour after recovering. *p = 0.0002. (−ATR, −light vs +ATR, +light p < 0.0001, –ATR, +light vs. +ATR, +light p = <0.0001. n = 3 plates each condition). Statistics are one-way ANOVA with Tukey’s post hoc test. Bars are means ± SEM.

### mtON affects whole-animal energy sensing

We next asked whether mtON could affect metabolic signaling. One way organisms sense energy availability and preserve energy homeostasis is through the AMP-activated protein kinase (AMPK) (Hardie, Ross et al., 2012). In *C. elegans* the *aak-2* gene encodes the catalytic subunit ortholog of mammalian AMPKα2. Mutation of *aak-2* has well-characterized phenotypic outputs linked to energy availability, and serves as a regulator of whole-organism energy sensing (Apfeld, O’Connor et al., 2004, Cunningham, Bouagnon et al., 2014). We hypothesized that mtON activity would signal energy availability and decrease phosphorylation of AMPK, its activated state. As such, we exposed animals expressing mtON to light in the absence of food, where AMPK should be phosphorylated, and immunoblotted against phosphorylated AMPK. We found that removal from food increases AMPK phosphorylation as expected, and mtON activation prevented this phosphorylation (Fig. 4A). In *C. elegans,* the AMPK homologue AAK-2 regulates a behavioral response to food availability (Lee, Cho et al., 2008), where in the absence of food animals will increase locomotion to search for food. Based on our phosphorylation results, we hypothesized that mtON activation would attenuate the energy deficit signal for animals off food, suppressing their movement, and thus mimicking the *aak-2* loss-of-function phenotype. We first confirmed that *aak-2* loss-of-function mutant animals had decreased locomotion in the absence of food compared to wild type (Fig. 4B). In response to acute mtON activation in wild type background, we observed decreased locomotion in the absence of food (Fig. 4C). This effect was reversed using the AMPK activator 5-aminoimidizole-4-carboxamide ribonucleotide (AICAR), indicating the mtON suppression of locomotion was rescued by AMPK activity (Fig. 4C). These data suggest that mtON activity can modify downstream metabolic signaling. As changing the PMF affects many aspects of metabolism that are able to activate AMPK, the exact mechanism of AMPK activation in our experiments is unclear. For example, AMPK is activated by increased energy demand, but also redox signaling (Garcia & Shaw, 2017), and calcium signaling (Hardie, 2011). Upon further characterization in these contexts mtON could be used to modulate the many cellular processes that AMPK regulates (Hinchy, Gruszczyk et al., 2018, Jeon, 2016, Mihaylova & Shaw, 2011, Trewin, Berry et al., 2018b).

**Figure 4.**
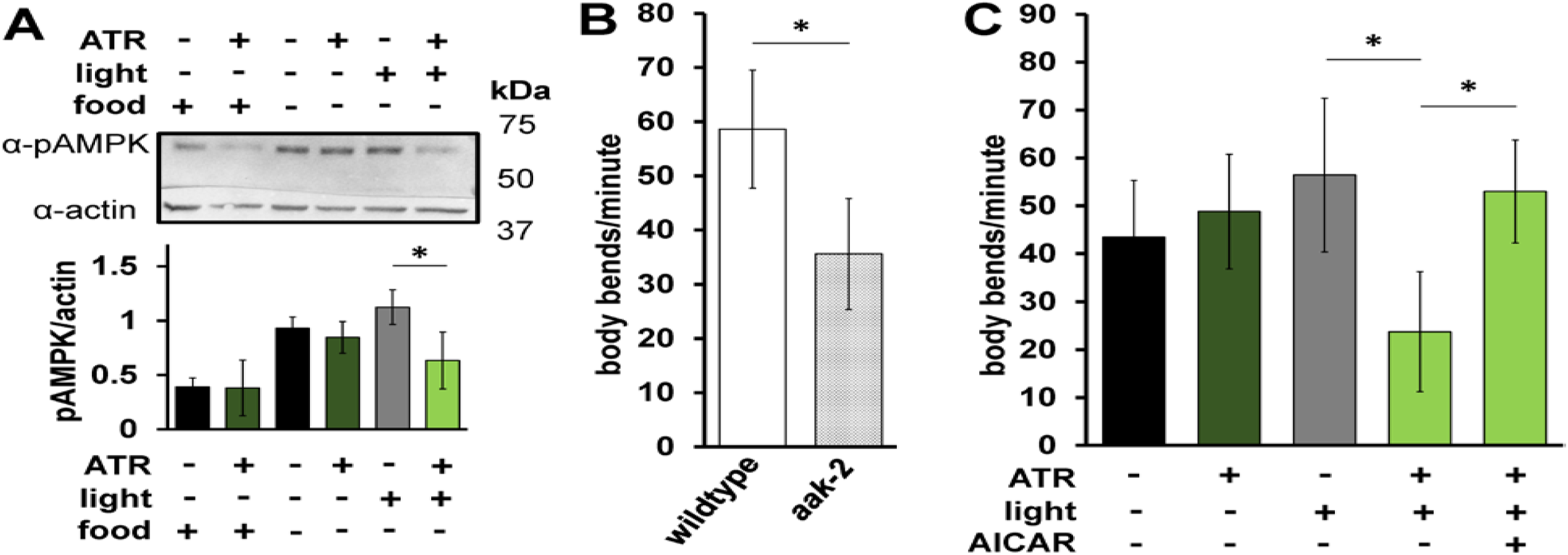
mtON affects whole-animal energy sensing. **A** Top: Immunoblot assessing the effect of mtON activation on AMPK phosphorylation status. Top bands (∼62 kDa) are phosphorylated AMPK signal, and bottom bands (∼43 kDa) are actin signal. Image is from the same membrane cut to separately probe for phosphorylated AMPK (pAMPK) and actin. Bottom: Densitometry analysis showing decreased pAMPK to actin ratio, as there is no known antibody directed against total AMPK in *C. elegans.* Phosphorylation increases in the absence of food, but low phosphorylation is preserved when mtON is activated, two-way ANOVA with Sidak’s test for multiple comparisons, *p = 0.0203, n = 3-4 biological replicates. **B** Locomotion was assessed by counting body bends per minute. Wild type animals were compared to *aak-2(ok524)* mutant animals. 2-sample, 2-tailed unpaired t-test *p < 0.0001, wild type n = 35, *aak-2* n = 39 animals across at least 3 days. Bars are means ± standard deviation. **C** Locomotion in response to mtON activation. Illumination was continuous throughout body bends measurement (see methods). For AAK-2 activation, animals were exposed to 1 mM AICAR for 4 hours before body bends measurement. One-way ANOVA with Tukey’s post hoc test, −ATR, +light vs +ATR, +light *p < 0.0001, +ATR,+light vs AICAR *p < 0.0001. n = (in order, left to right) 36, 39, 37, 46, 36 animals across at least 3 days. Bars are means ± standard deviation.

### mtON inhibits hypoxia-adaptation

We next tested if mtON could impact stress resistance. Hypoxia and reoxygenation (HR) is a pathologic insult that involves changes in the PMF that can contribute to injury and survival (Chouchani, Pell et al., 2014, Michiels, 2004, Murphy & Hartley, 2018, Sanderson, Reynolds et al., 2013, Solaini et al., 2010, Xu, Wang et al., 2001, Yang et al., 2018) depending on the context and degree of (de)polarization. In addition, the phenomenon of preconditioning (PC) is effectively modeled with hypoxia in *C. elegans* (Dasgupta, Patel et al., 2007, Hayakawa, Kato et al., 2011, Jia & Crowder, 2008, Pena et al., 2016, Wojtovich, DiStefano et al., 2012a), where a short period of hypoxia protects against a later pathologic exposure (Fig. 5B). A decreased PMF before HR also mediates a protective effect (Brennan, Berry et al., 2006, Brennan, Southworth et al., 2006, Ozcan, Palmeri et al., 2013), but mechanisms are debated (Shabalina & Nedergaard, 2011), as understanding is complicated by lack of temporal control when using pharmacologic approaches. In support of this, the protonophore FCCP, which can dissipate the PMF, protected wild type animals against HR (Fig. 5C), suggesting hypoxia resistance may be mediated by a decreased PMF. We combined these lines of evidence and hypothesized that increasing the PMF during PC would reverse the protection against hypoxia afforded by PC. To test this hypothesis we activated mtON selectively during PC (Fig. 5A). To quantify protection, we subtracted the percent survival after pathologic hypoxia from the percent survival after an intervention, giving the percent alive above baseline (Fig. 5C). Further highlighting the importance of our control conditions, we found that light on its own was as protective as PC in mtON expressing animals, and the combination of light and PC was additive (Fig. 5D). However, activation of mtON during PC decreased the protection (Fig. 5D), supporting our hypothesis that some of the protective effect of PC is mediated by decreasing the PMF. In effect, mtON counteracts a PC-induced decrease in the PMF by restoring it to normal levels. These data suggest that loss of PMF during PC is necessary for hypoxic adaptation. The FCCP data corroborate our mtON findings, and demonstrate the sufficiency of PMF loss to protect against hypoxia. Using mtON only during PC to boost the PMF provided temporal control that had not been tested before, allowing us to determine when PMF changes have physiologic effects in HR.

**Figure 5.**
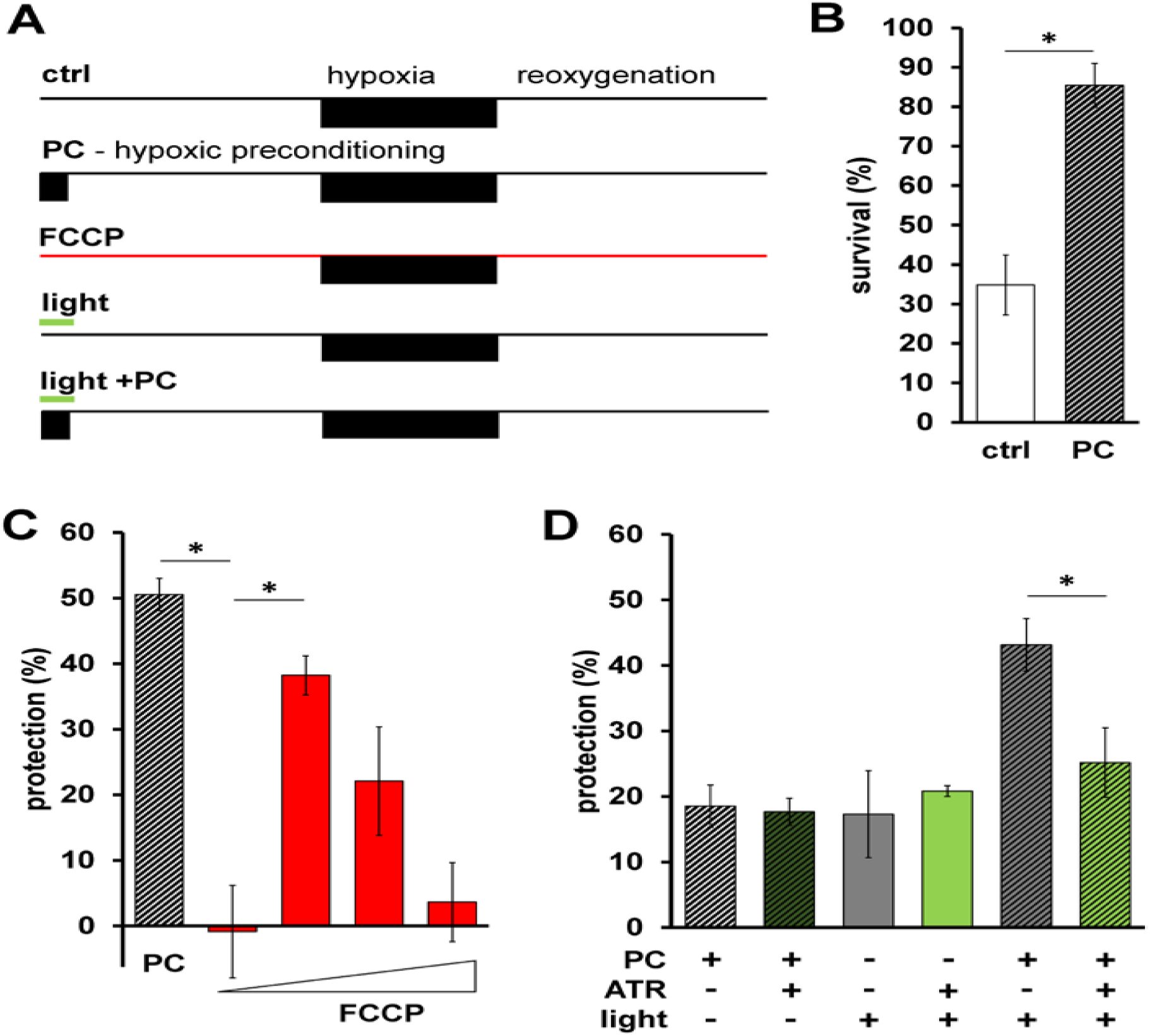
mtON inhibits hypoxia-adaptation. **A** Schematic of HR experiments. Top shows control hypoxia and reoxygenation. Below is the hypoxic preconditioning (PC) protocol, survival for these two timelines shown in panel B. Third from the top shows FCCP treatment protocol, calculated protection shown in panel C. Bottom two show light treatment protocol + and - PC, calculated protection data from these shown in panel D. **B** Survival after HR in day 1 adult animals is shown with hypoxic preconditioning (PC, represented by diagonal stripes) 2-sample 2-tailed unpaired t-test, ctrl vs. PC *p = 0.0035, n = 4, each averaged from 3 technical replicates. Bars are means ± SEM. **C** Protection (%) is percent survival minus percent survival of control condition. PC data calculated from panel B. FCCP final concentrations 0.001, 0.01, 0.1, 1 nM. One-way ANOVA with Tukey’s post hoc test, PC vs. 0.001 nM FCCP, p = 0.0012, 0.001 nM FCCP vs 0.01 FCCP, p = 0.0142, n = 3 - 4 independent experiments, each averaged from 3 technical replicates. Bars are means ± SEM. **D** Illumination was continuous throughout PC alone, control illumination was for the same duration under normoxic conditions (see methods & panel A). Two-way ANOVA comparing +ATR vs −ATR in each group. *p = 0.016, n = (in order, left to right) 11,11,4,4,6,6 independent experiments, each averaged from 3 technical replicates. Bars are means ± SEM.

Our findings that protection afforded by PC relies on the transient loss of PMF suggest that a decreased PMF is sufficient to elicit stress resistance at a later period. This implies that interventions at the level of the PMF alone can impact cellular responses to stress. Combining the mtON approach with tissue-specific gene promoters, precise spatiotemporal control could be achieved in discerning the effects of mitochondrial function across tissues in whole organisms. Taken together, mtON appears to be a useful tool to enable discrimination of cause and effect in complex (patho)physiologic contexts that involve modest changes in mitochondrial function and metabolism. Defining when changes in the PMF are adaptive and when they are detrimental may advance our understanding of many pathologies and inform novel therapeutic strategies that target mitochondrial function. Our findings from this approach suggest the PMF is a keystone of metabolism that senses cellular stress and elicits appropriate adaptive responses to maintain homeostasis.

## MATERIALS & METHODS

### Molecular biology

The light-activated proton pump from *Leptosphaeria maculans* (Mac) fused to eGFP was amplified from plasmid DNA pFCK-Mac-GFP, a gift from Edward Boyden (Addgene plasmid #22223 (Chow et al., 2010)) (forward amplification primer: ACACCTGCAGGCTTGATCGTGGACCAGTTCGA, reverse amplification primer: CACAGCGGCCGCTTACTTGTACAGCTCGTCCA). The N-terminal 187 amino acids of the *Immt1* gene was amplified by PCR from mouse cDNA (forward amplification primer: ACAACCGGTAAAAATGCTGCGGGCCTGTCAGTT, reverse amplification primer: CACCCTGCAGGTTCCTCTGTGGTTTCAGACG). The ubiquitously expressed gene promoter P*eft-3* (also known as P*eef-1A.1*) was amplified by PCR from pDD162 (forward amplification primer: AACAAAGCTTGCACCTTTGGTCTTTTA, reverse amplification primer: ACATCTAGAGAGCAAAGTGTTTCCCA). The body wall muscle promoter P*myo-3* was PCR amplified from pDJ16 (forward amplification primer: ACAGCTAGCTGTGTGTGATTGCT, reverse amplification primer: ACAACCGGTGCGGCAATTCTAGATGG). PCR fragments were ligated into pFH6.II (pPD95.81 with a modified multi-cloning site) for *C. elegans* expression using restriction digest cloning. Resulting plasmids were pBJB20 (P*eft-3*::IMMT1(N-terminal 187 amino acids)::Mac::GFP), pBJB16 (P*myo-3*::IMMT1(N-terminal 187 amino acids)::Mac::GFP), Sanger sequencing was used to confirm plasmid sequences (Eurofins Genomics). Animals were transformed by plasmid DNA microinjection with *pha-1(+)* selection in *a pha-1(e2123ts)* temperature-sensitive mutant strain, where transgenic animals were selected for growth at 20° C.

### *C. elegans* strains growth and maintenance

All animals were maintained at 20° C on nematode growth medium (NGM) seeded with OP50 *E. Coli.* Young adult hermaphrodite animals were used for all experiments. Transgenic strains were generated by plasmid DNA microinjection as described (Mello, Kramer et al., 1991). For a complete strain list see Table 1. Where indicated, OP50 was supplemented with all trans-Retinal to a final concentration of 100 μM on seeded NGM plates. Animals were cultured on ATR-containing plates for at least one generation.

**Table 1.**
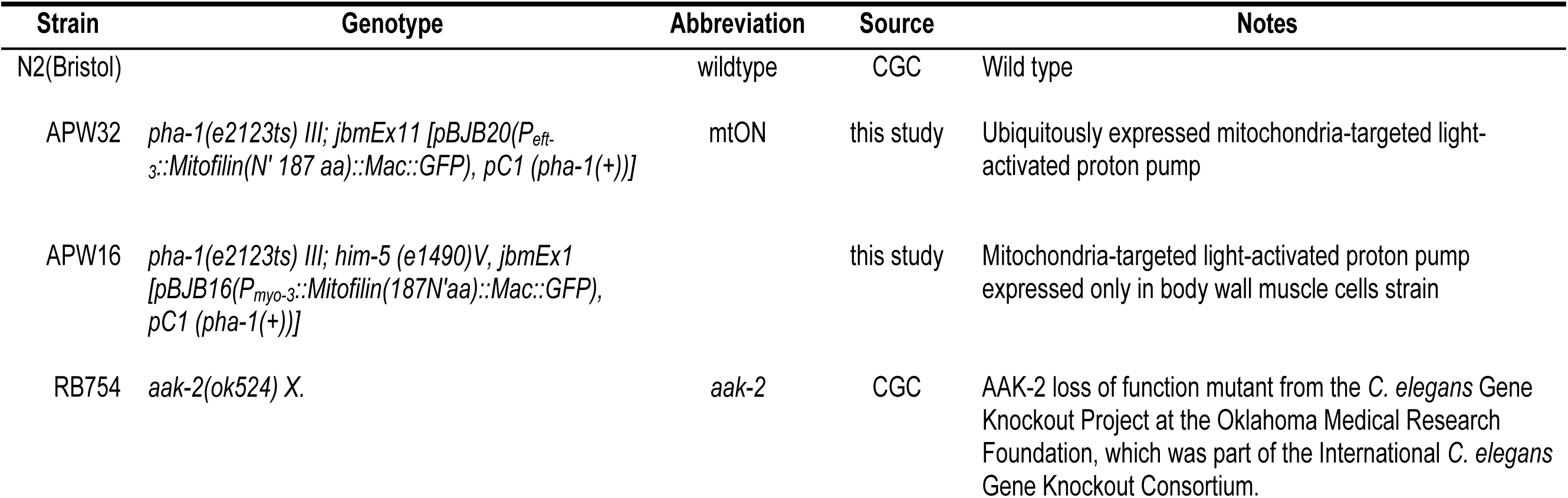
*C. elegans* strains. Some strains were provided by the C. elegans Genetics Center (CGC).

### Fluorescence microscopy

Images were taken on a FV1000 Olympus laser scanning confocal microscope using a 60x oil objective (Olympus, N.A. 1.42). Diode laser illumination was at 561 nm for red fluorescence and 488 nm for green fluorescence. Where indicated, animals were stained with 10 µM MitoTracker^TM^ Red CMXRos for 4 hours. MitoTracker^TM^ stain was dissolved in DMSO, diluted in M9 media (22 mM KH_2_PO_4,_ 42 mM Na_2_HPO_4,_ 86 mM NaCl, 1 mM MgSO_4_, pH 7) and added to OP50 seeded NGM plates (DMSO < 0.02% final) and allowed to dry. Line scan pixel intensity was performed using ImageJ software.

### Mitochondria isolation

*C. elegans* mitochondria were isolated from day 1 adults as previously described (Wojtovich et al., 2012a) using differential centrifugation in mannitol and sucrose-based media. Animals from three 15 cm culture plates were transferred into 50 mL of M9 media in a conical tube and allowed to settle by gravity on ice. Pelleted animals were rinsed with ice-cold M9 twice, and once with ice-cold mitochondrial isolation media (220 mM mannitol, 70 mM sucrose, 5 mM MOPS, 2 mM EGTA, pH 7.4) with 0.04% BSA. After settling by gravity, supernatant was removed and worms were transferred to an ice-cold mortar containing ∼2 g of pure sea sand per 1 mL of animals. Animals were ground with an ice-cold pestle for 1 minute and extracted from the sand using mitochondrial isolation media and transferred to a 10 mL conical tube. The suspension was then transferred to an ice-cold glass Dounce homogenizer and homogenized with 40 strokes. The homogenate was centrifuged at 600 g for 5 minutes. Supernatant was transferred to a new tube and centrifuged at 700 g for 10 minutes. The pellet was resuspended in 1 mL of mitochondrial isolation media without BSA, which was centrifuged at 7000 g for 5 minutes. The pellet was finally resuspended in 50 μL of mitochondrial isolation media without BSA. Protein concentration was quantified using the Folin-phenol method.

### Light sources

Illumination sources included a 580 nm Quantum SpectraLife LED Hybrid lamp by Quantum Devices, Barneveld WI, USA (abbreviated Quantum LED), a 540-600 nm GYX module, X-Cite LED1 by Excelitas, Waltham MA, USA (abbreviated XCite LED), and a 540-580 nm excitation filter MVX10 Fluorescence MacroZoom dissecting microscope by Olympus (abbreviated MVX).

Light intensities are indicated for each experimental condition and were determined with a calibrated thermopile detector (818P-010-12, Newport Corporation, Irvine, CA) and optical power meter (1916-R, Newport Corporation).

### Immunoblotting

Synchronized young adult animals were exposed to 1 Hz light (Quantum LED, 0.02 mW/mm^2^) for 4 hours, and immediately harvested with ice-cold M9 media and centrifuged at 1,000 x g for 1 minute. Animals were ground by plastic pestle disruption in lysis buffer (20mM Tris-HC, 100 mM NaCl, 1mM EDTA, 1mM DTT, 10% glycerol, 0.1% SDS, pH 7.6, 1X Halt^TM^ protease inhibitor cocktail, Thermo78429) and diluted 1:1 in sample loading buffer (100 mM Tris HCl, 10% v/v glycerol, 10% SDS, 0.2% w/v bromophenol blue, 2% v/v β-mercaptoethanol). These samples were heated at 95 °C for 5 minutes. Isolated mitochondrial samples were prepared as described above and diluted 1:1 in sample loading buffer with 1% SDS. 30 μg samples were loaded to 7.5% polyacrylamide gels and separated by SDS-PAGE. Proteins were transferred to nitrocellulose membranes and blocked in 5% nonfat milk/TBST (50 mM Tris, 150 mM NaCl, 0.05% Tween 20, pH 8.0) for 1 h at room temperature. Membranes were incubated at 4°C in primary antibodies diluted 1:1000 in 5% bovine serum albumin: anti-GFP (ClonTech Living Colours #ab632375), anti-HSP60 (Department of Biology, Iowa City, IA 52242. Developmental Studies Hybridoma Bank, University of Iowa Department of Biology, Iowa City, IA 52242), (Cell Signaling, #4188), anti-Actin (Abcam #ab14128), and 1:10,000 anti-phospho-AMPKα). Membranes were washed in TBST and incubated in horseradish peroxidase-conjugated secondary antibodies: anti-rabbit IgG (Cell Signaling #7074S) or anti-mouse IgG (Thermo Scientific #32430, lot #RF234708) for 1 hour at room temperature. Proteins were visualized using ECL (Clarity Western ECL Substrate, Bio Rad) by chemiluminescence (ChemiDoc, Bio Rad). Densitometry was performed using Image Lab software (version 5.2.1).

### Mitochondrial membrane potential measurement

Isolated mitochondria at 0.5 mg/mL were stirred in mitochondrial respiration buffer (MRB: 120 mM KCl, 25 mM sucrose, 5 mM MgCl_2_, 5 mM KH_2_PO_4_, 1 mM EGTA, 10 mM HEPES, 1 mg/mL FF-BSA, pH 7.35) at 25° C in the presence of 2 μM rotenone, and 5 mM succinate where indicated. 300 nM tetramethylrhodamine, ethyl ester (TMRE, ThermoFisher, T669) was added to observe mitochondrial membrane potential in quench mode. Under quenching conditions TMRE fluorescence is low in the presence of a Δψ_m_. Upon addition of a protonophore (e.g. FCCP), TMRE will exit mitochondria and dequench and increase total fluorescence (Chouchani et al., 2014, Perry, Norman et al., 2011). TMRE signal was measured by Cary Eclipse Fluorescence Spectrophotometer (Agilent Technologies) using a 335-620 nm excitation filter and a 550-1100 nm emission. Illumination was performed continuously throughout all measurements (555 nm, 0.0016 mW/mm^2^). Increasing illumination time exposed mitochondria to more photons (calculated as fluence, J/cm^2^). After stable baseline measurements with or without succinate, 2 μM FCCP was added to completely depolarize mitochondria. The average fluorescence intensity after addition of 2 μM FCCP (maximum fluorescence in quench mode) was subtracted from the starting test condition to give a change in fluorescence corresponding to changes in Δψ_m_ (ΔF for conditions without succinate, and ΔF_max_ for conditions with succinate). To represent polarization of the Δψ_m_, we used the ratio of change in fluorescence (ΔF, FCCP fluorescence minus experimental fluorescence) to the maximum change in fluorescence generated by succinate-driven Δψ_m_ (ΔF_max_).

### Mitochondrial matrix pH measurement

The ratiometric pH indicator BCECF-AM (ThermoFisher, B1170) was used to measure pH changes in the mitochondrial matrix (Aldakkak, Stowe et al., 2010) in response to succinate respiration or mtON activation. Isolated mitochondria (∼200 μL per isolation) were incubated at room temperature with 50 μM BCECF-AM for 10 minutes with periodic mixing. Mitochondria were then pelleted at 7000 g for 5 minutes at 4° C, isolation media replaced and pelleted again to remove extramitochondrial BCECF-AM. Isolated mitochondrial suspensions were then assayed under the same conditions as in the mitochondrial membrane potential measurements described above. Ratiometric fluorescent signal was measured by Cary Eclipse Fluorescence Spectrophotometer (Agilent Technologies) using 440 and 490 nm excitation wavelengths and 545 nm emission. The fluorescence intensity ratio at 545 nm of 490/440 nm excitation wavelengths was used to represent pH changes in the mitochondrial matrix. Light treatment was 0.16 J/cm^2^ (XCite LED, 0.02 mW/mm^2^), and 2 μM FCCP was used at the end of each trace to establish baseline signal.

### ATP measurement

Relative ATP levels were determined in isolated mitochondria using a luciferase bioluminescence kit according to manufacturer’s instructions (Invitrogen^TM^ Molecular Probes^TM^, A22066). Mitochondria were stirred in MRB at 0.5 mg/mL with 1 mg/mL fat free BSA, 600 μM ADP, 2 μM rotenone. 5 mM succinate for a control for maximum ATP level, and 0.001 mg/mL oligomycin A was used as a zero ATP synthesis control. Mitochondrial suspensions were immediately frozen with liquid nitrogen after 1, 5, or 10 minutes light exposure (XCite LED, 0.02 mW/mm^2^). Samples were then thawed on ice, centrifuged at 14,800 g, and supernatant was collected and run at 1:100 dilution in MRB in the luminescence assay. Oligomycin A control values were subtracted from experimental reads, and data were then normalized to luminescent signal from succinate control samples (complete ADP conversion confirmed by monitoring O_2_ consumption rate transitions).

### Mitochondrial O_2_ consumption

O_2_ consumption was measured using a Clark-type O_2_ electrode (S1 electrode disc, DW2/2 electrode chamber and Oxy-Lab control unit, Hansatech Instruments, Norfolk UK) at 25° C. Isolated mitochondria were stirred in MRB at 1 mg/mL with 1 mg/mL fat free BSA. Substrates and inhibitors were added by syringe port (100 μM ADP, 2 μM rotenone, 5 mM succinate). Given excess succinate as substrate for ETC respiration, we measured the amount of O_2_ required to convert 50 nmol of ADP to ATP and established a baseline for comparison. To test mtON activity, we illuminated mitochondria as in the ATP measurement, for 1, 5, or 10 minutes in the presence of ADP without succinate to allow for mtON conversion of ADP to ATP. Any remaining ADP was then converted using O_2_-dependent ETC respiration upon succinate addition (Fig. EV4C). Slopes were calculated from plots of O_2_ concentration versus time to give rates of O_2_ consumption during ADP respiration, ADP+succinate respiration, and respiration after ADP had been entirely consumed. The intersections of these three rates were used to calculate total amount of O_2_ consumed during ADP+succinate respiration. Light activation of mitochondria (XCite LED, 0.02 mW/mm^2^) during ADP respiration alone was carried out for differing lengths of time to test the ability of mtON to drive ADP consumption before succinate was added. Dark control was 10 minutes of no illumination before addition of succinate.

### ETC inhibitor assays

Experiments were performed on at least three separate days, using 15-100 young adult animals per plate. Seeded plates were supplemented with rotenone (50 μM final concentration), or antimycin A (50 μM final concentration) 24 hours before animals were transferred onto them. For azide toxicity animals were placed in M9 buffer with 250 mM azide. Control plates were kept in the dark and experimental plates were exposed to 1 Hz light (Quantum LED, 0.02 mW/mm^2^) for the duration of the experiment. For rotenone, surviving animals were scored after 5 hours. For antimycin A, animals were scored 16 hours after exposure. For azide, animals were exposed for one hour in M9 and allowed to recover for 1 hour on a seeded culture plate, then survival was scored. Azide experimental plates were exposed to light (XCite LED, 0.19 mW/mm^2^) for the duration of azide treatment and recovery. For all toxins, animals that were moving or those that moved in response to a light touch to the head were scored as alive.

### Locomotion assay

Locomotion was scored by counting the number of body bends in 15 seconds immediately after being transferred off OP50 food (Sawin, Ranganathan et al., 2000) (n = 36-46 animals scored on at least 2 separate days). 1 body bend was scored as a deflection of direction of motion of the posterior pharyngeal bulb (Tsalik & Hobert, 2003). For AMPK activation, animals were placed on plates containing 1 mM AICAR 4 hours before counting body bends. AICAR was dissolved in M9 buffer, added directly onto OP50 seeded plates and allowed to dry. Illumination was continuous through measurements (MVX, 0.265 mW/mm^2^).

### Hypoxia and reoxygenation

Experiments were carried out using a hypoxic chamber (Coy Lab Products, 5%/95% H_2_/N_2_ gas, palladium catalyst) at 26° C with 50-100 animals per plate. O_2_ was less than 0.01%. Hypoxic preconditioning (PC) duration was 4 hours, and control animals were incubated at 26° C in room air for the same time. 1 Hz illumination (Quantum LED, 0.02 mW/mm^2^) was carried out during PC period alone. Hypoxic exposure was for 18.5 hours, 21 hours after PC. 24 hours after hypoxia exposure, animals that were moving or those that moved in response to a light touch to the head were scored as alive. Data from days where PC was at least 15% effective for both +ATR and −ATR were used. Animals supplemented with ATR were allowed to lay eggs onto plates with no ATR that were subsequently used for HR experiments to minimize confounding effects of ATR. ATR from the parent animal was sufficient to provide active mtON in progeny as tested by the azide toxin assay described above. Only progeny from parents supplemented with ATR were significantly protected against azide upon illumination (tested by One-way ANOVA, −ATR, −light [17.3% survival] vs. +ATR, +light [66.8% survival] p = 0.0054, −ATR, +light [32.5% survival] vs. +ATR, +light p = 0.0395, +ATR, −light [27.5% survival] vs. +ATR, +light p = 0.0199, DF = 11, F = 8.94). PC experiments were represented as protection (%), where baseline survival was subtracted to correct for protection from ATR (−ATR survival: 34.4 ± 14.4%, +ATR survival: 49.7 ± 20.3%, 2 sample 2-tailed paired t-test, p = 0.033, n = 12 independent experiments). FCCP dissolved in ethanol was deposited onto seeded NGM plates to the final concentrations indicated in the figure legend (Fig. 5C).

### Statistics

One-way ANOVA was used when comparing only our four experimental conditions (Fig. EV2). Two-way ANOVA was used when comparing our conditions with other variables, such as +/- succinate. Shapiro-Wilkes normality tests were used to determine whether parametric or non-parametric tests should be used. See figure legends for detailed statistical information and post-hoc tests.

## DATA AVAILABILITY

All data are presented in the manuscript, and raw files will be made available upon request.

## ACKNOWLEDGEMENTS

Work in the laboratory of A.P.W. is supported by a grant from National Institutes of Health (R01 NS092558) and institutional funds from the University of Rochester; B.J.B. is supported by an American Heart Association Predoctoral Fellowship (18PRE33990054) and an Institutional Ruth L. Kirschstein National Research Service Award (NIH T32 GM068411). A.J.T current address: Institute for Physical Activity and Nutrition (IPAN), Deakin University, Burwood, Australia. We thank the members of the mitochondrial research groups at University of Rochester Medical Center for helpful discussions, suggestions, and guidance. Some strains were provided by the CGC, which is funded by NIH Office of Research Infrastructure Programs (P40 OD010440).

## AUTHOR CONTRIBUTIONS

B.J.B. and A.P.W. designed the research. B.J.B., A.J.T., A.S.M., A.B. and A.M.A. performed experiments. M.K. provided critical review of the results and manuscript. B.J.B. analyzed the data and wrote the manuscript. All authors approved the final version of the manuscript.

## CONFLICT OF INTEREST

The authors declare that they have no conflict of interest.

## Expanded view figures

**Figure EV1.**
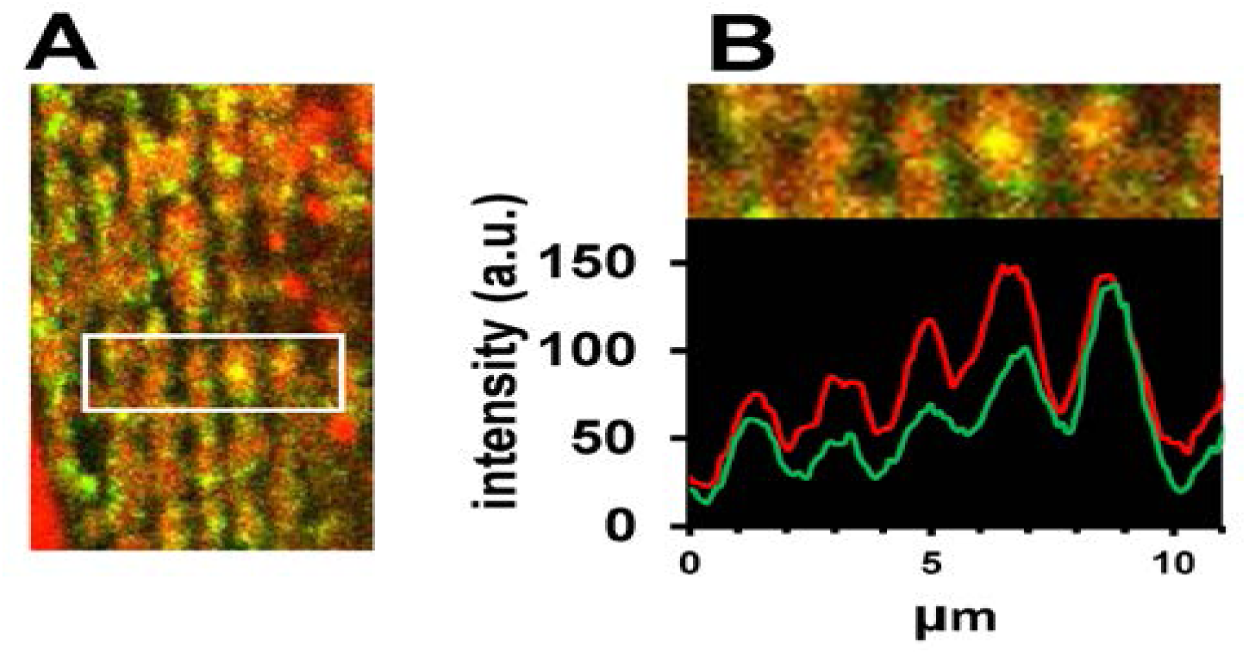
mtON is targeted to mitochondria. **A** mtON tagged with GFP expressed in body wall muscle overlaps with MitoTracker^TM^ Red CMXRos stained mitochondria. **B** Profile plot of a selected region demonstrating the overlapping signal intensity of GFP and MitoTracker^TM^ Red CMXRos determined using ImageJ software.

**Figure EV2.**
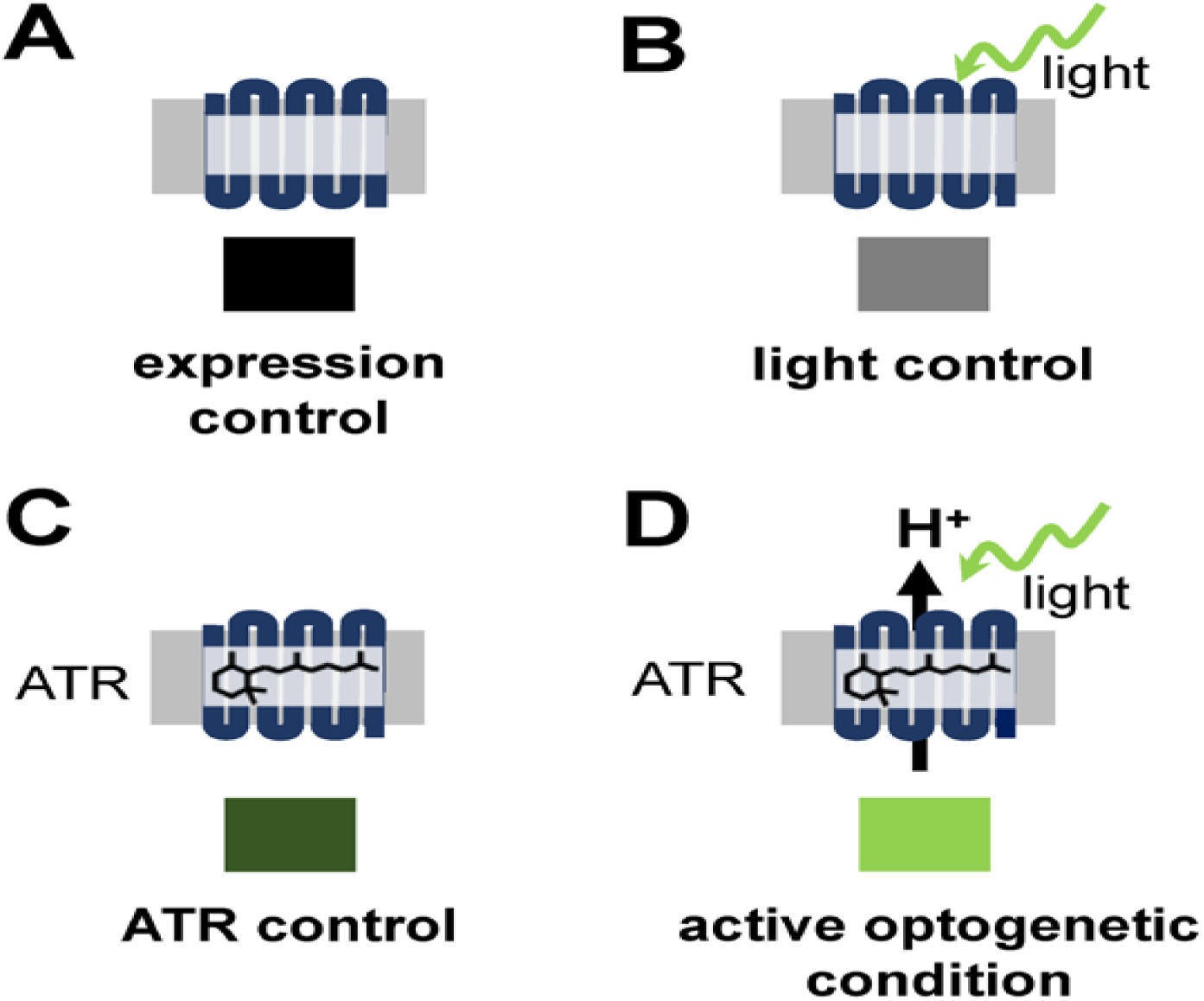
Schematic of experimental conditions. **A** Expression control: baseline condition where animals express mtON, but have not been supplemented with ATR or exposed to light. This condition is represented by the color black throughout. **B** Light control: condition exposed to light where mtON is illuminated but not supplemented with ATR, resulting in an inactive proton pump. This condition is represented by the color gray throughout. **C** ATR control: condition supplemented with ATR but not exposed to light, where proton pumping is possible but no light activation has occurred. This condition is represented by dark green throughout. **D** Active optogenetic condition: supplemented with ATR and exposed to light, where proton pumping activity is expected. This condition is represented by bright green throughout.

**Figure EV3.**
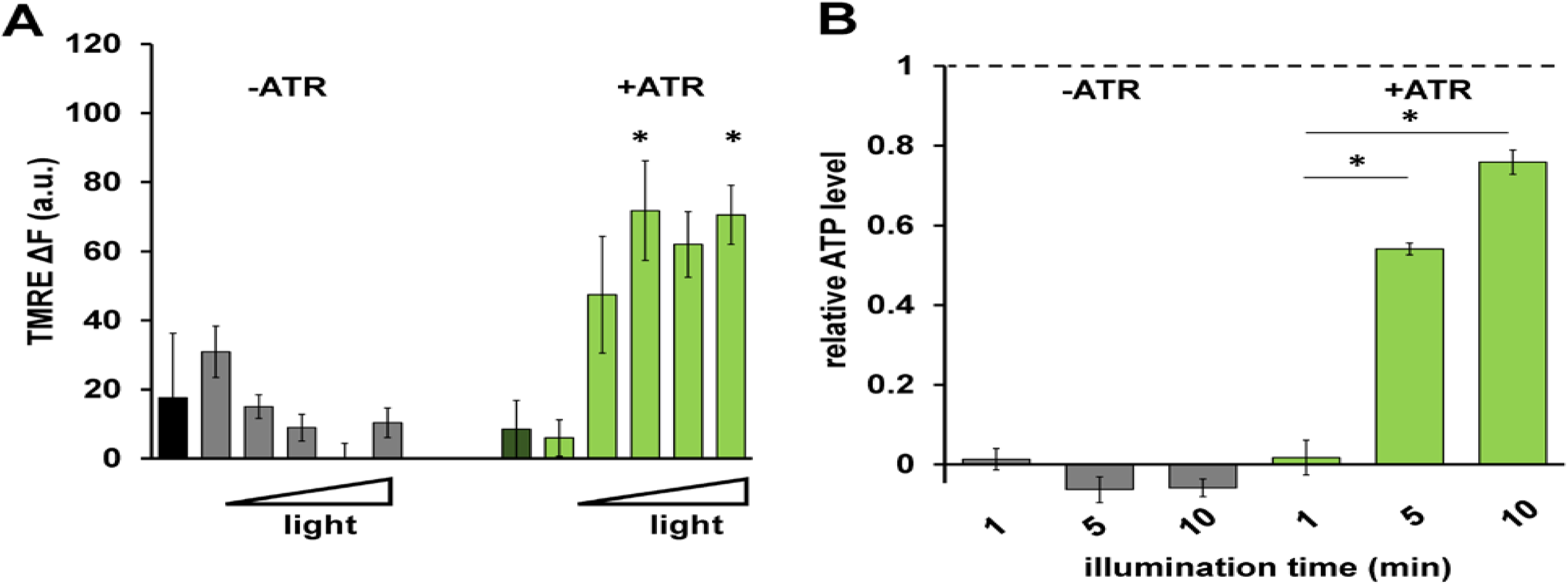
mtON increases the PMF and ATP synthesis light dose-dependently. **A** Quantification of change in TMRE fluorescence with increasing light dose (increasing fluence from left to right: 0, 0.016, 0.08, 0.096, 0.16, 0.24 J/cm^2^). Two-way ANOVA with Sidak’s multiple comparisons test, +ATR: 0 vs. 0.096 J/cm^2^ *p = 0.0232, 0 vs. 0.24 J/cm^2^ *p = 0.0149, 0.016 vs. 0.096 J/cm^2^ *p = 0.0162, 0.016 vs. 0.16 J/cm^2^ *p = 0.0376, 0.016 vs. 0.24 J/cm^2^ *p = 0.0101. n = 3 - 6 mitochondrial isolations. **B** ATP levels in response to increasing illumination normalized to maximum ATP synthesis given by succinate respiration (dotted line). 10 minute illumination time from Fig. 2E. Two-way ANOVA with Sidak’s multiple comparison test, −ATR: succinate vs. 1, 5 and 10 minutes light *p < 0.0001. +ATR: succinate vs. 1 minute light, *p < 0.0001, 1 minute light vs. 5 minutes light *p = 0.0098, 1 minute light vs. 10 minutes light *p = 0.0001. n = 3 - 7 light treatments across 2 mitochondrial isolations.

**Figure EV4.**
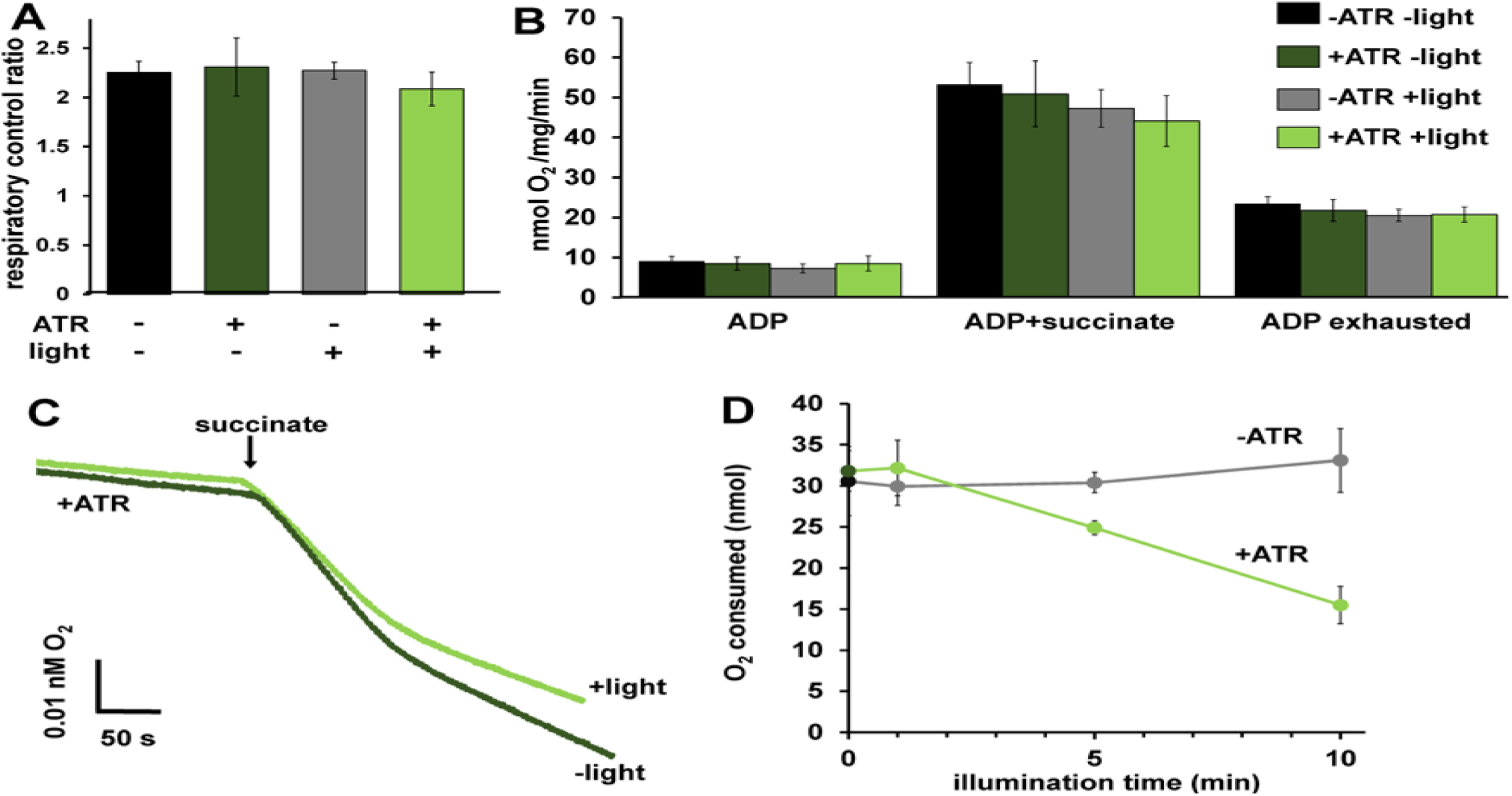
mtON effects on respiration. **A** Respiratory control ratios (indicative of ability of isolated mitochondria to respond to energy demand) were calculated by dividing the rate of O_2_ consumption with ADP and succinate by the rate after ADP was depleted. One-way ANOVA with Tukey’s post hoc test, p = 0.72, n = 6 mitochondrial preparations. Bars are means ± SEM. **B** O_2_ consumption rates under different states of respiration comparing the control conditions. One-way ANOVA with Tukey’s post hoc tests performed for each respiration state. ADP rate p = 0.66, Succinate rate p = 0.61. ADP exhausted rate p = 0.83. Bars are means ± SEM. **C)** Representative traces depicting O_2_ consumption rate. +/- light for mitochondria with ATR present. Light exposure was 10 minutes before the addition of succinate. Traces show an initial rapid depletion of ADP before transitioning to a lower rate of O_2_ consumption. **C** Activation of mtON decreases the amount of O_2_ required to consume 50 nmoles ADP light dose-dependently. Isolated mitochondria were exposed to light for the indicated time and then succinate was added (example traces in panel c). Dark and 10 minute illumination data was used for analysis in Fig. 2F. Linear regression showed a negative relationship between O_2_ required to consume ADP and illumination time in mitochondria from animals with mtON that were supplemented with ATR, R^2^ = 0.98 p = 0.007, n = 6 mitochondrial preparations. Linear regression shows no relationship between O_2_ required to consume ADP and illumination in mitochondria from animals with mtON not supplemented with ATR, R^2^ = 0.756 p = 0.130, dark n = 5, 1 min light n = 5, 5 min light n = 6, 10 min light n = 6. Data are means ± SEM.

**Figure EV5.**
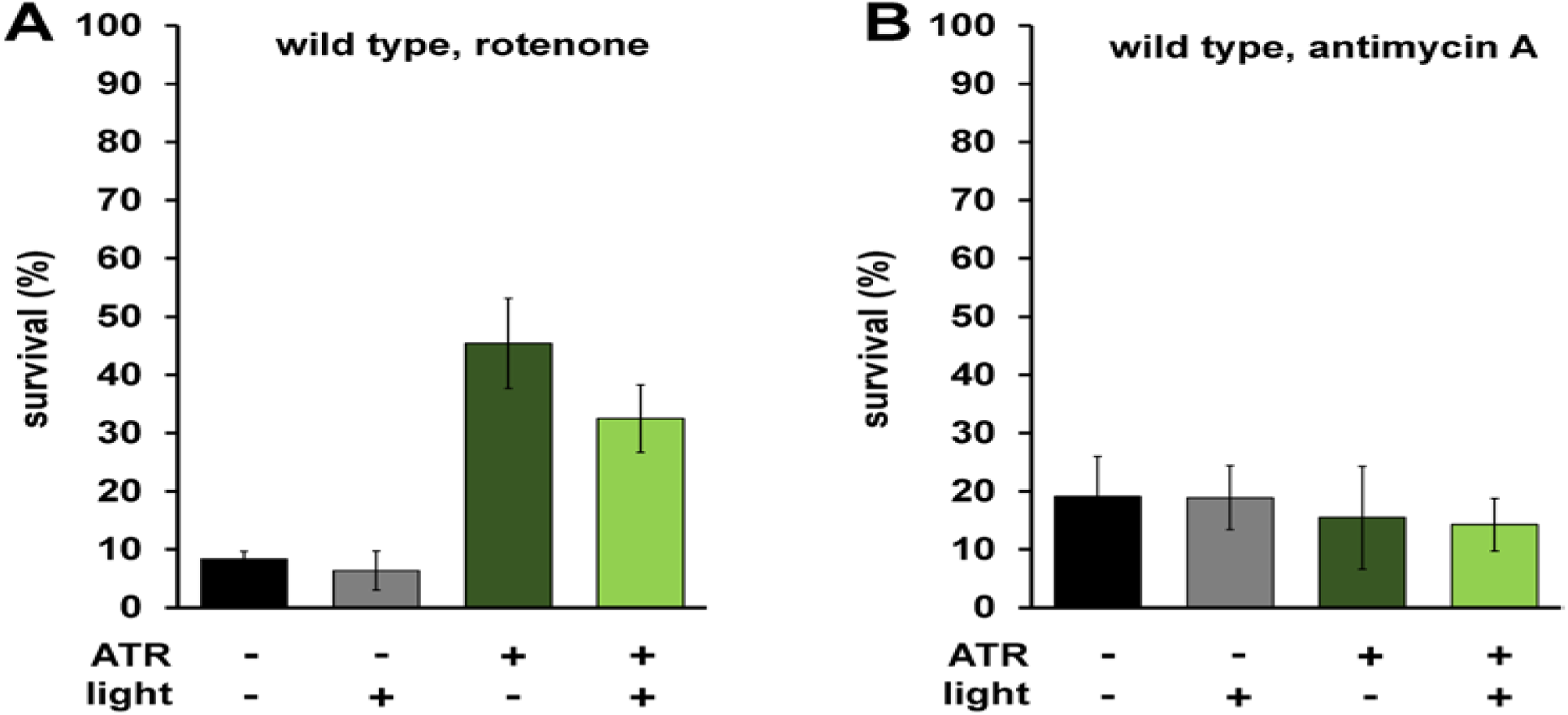
ATR and light have no effect in wild type survival. **A** Same experimental conditions from Fig. 3A, here testing wild type animals. No difference in survival within ATR conditions, suggesting the interaction between ATR and light has no effect on its own. One-way ANOVA with Tukey’s multiple comparison test. Again, ATR alone was protective (−ATR, −light vs. +ATR, −light p = 0.0111.) **B** Same experimental conditions from Fig. 3B, here testing wild type animals, showing no difference in survival. One-way ANOVA with Tukey’s multiple comparison test.

